# The winning methods for predicting cellular position in the DREAM single cell transcriptomics challenge

**DOI:** 10.1101/2020.05.09.086397

**Authors:** Vu VH Pham, Xiaomei Li, Buu Truong, Thin Nguyen, Lin Liu, Jiuyong Li, Thuc D Le

## Abstract

**Motivation:** Predicting cell locations is important since with the understanding of cell locations, we may estimate the function of cells and their integration with the spatial environment. Thus, the DREAM Challenge on Single Cell Transcriptomics required participants to predict the locations of single cells in the Drosophila embryo using single cell transcriptomic data.

**Results:** We have developed over 50 pipelines by combining different ways of pre-processing the RNA-seq data, selecting the genes, predicting the cell locations, and validating predicted cell locations, resulting in the winning methods for two out of three sub-challenges in the competition. In this paper, we present an *R* package, *SCTCwhatateam*, which includes all the methods we developed and the *Shiny* web-application to facilitate the research on single cell spatial reconstruction. All the data and the example use cases are available in the Supplementary material.

**Availability:** The scripts of the package are available at https://github.com/thanhbuu04/SCTCwhatateam and the *Shiny* application is available at https://github.com/pvvhoang/SCTCwhatateam-ShinyApp

**Supplementary information:** Supplementary data are available at *Briefings in Bioinformatics* online.

## 1 Introduction

Single cell sequencing (scRNAseq) methods quantify the gene expression levels across thousands of cells of the same tissue, but the methods do not provide spatial information of the cells. The spatial information of cells is important in predicting functional roles of individual cells. Therefore, computational methods are required for reconstructing the spatial information [1, 6, 3] from single-cell RNA datasets and reference databases. The DREAM Challenge on Single Cell Transcriptomics used known spatial information of single cells in the early Drosophila embryo [4] as the ground-truth and required participants to develop computational methods to predict cell locations using only 60 genes (sub-challenge 1), 40 genes (sub-challenge 2), and 20 genes (sub-challenge 3) from the single cell RNA-seq dataset with 8924 genes of 1297 cells. As Karaiskos et al [4] had successfully reconstructed the cell locations using 84 marker genes, participants were given the expression levels of the 84 genes as the reference database in addition to the RNA-seq dataset.

We have developed methods to predict cell locations using less 84 genes (i.e. 60, 40 or 20 genes respectively). In our methods, we employ imputation to process missing data in the single cell RNA-seq dataset and select genes which are most unlikely to be predicted by other genes to be used in the prediction. In addition, we use the Matthews Correlation Coefficient score (MCC) [5] and the Local Outlier Factor (LOF) method [2] for cell location prediction.

Because of the excellent performance of our methods and pipelines in predicting cell locations, we won two (sub-challenges 1 and 2) out of the three sub-challenges. To make the winning methods accessible to researchers for spatial cell reconstruction using single cell RNA-seq data, we have implemented all of these methods as an *R* package, *SCTCwhatateam*. Moreover, we have built a *Shiny* web-application to allow users to upload their own datasets, customise their “recipe” by selecting methods in each component of the pipeline, and download the prediction results. As the winning methods of the challenge, we believe the package and the *Shiny* application will be useful for researchers to explore their data and/or to use our methods as the benchmark methods for testing their new methods.

## 2 Implementation and main functions

### 2.1 Overview

The *SCTCwhatateam* package contains three main components (steps) for predicting cell locations: data pre-processing, feature (gene) selection, and cell location prediction, as shown in Fig. 1. For each of the components, we have developed multiple methods. Using different methods in each component will result in different pipelines.

**Fig. 1.**
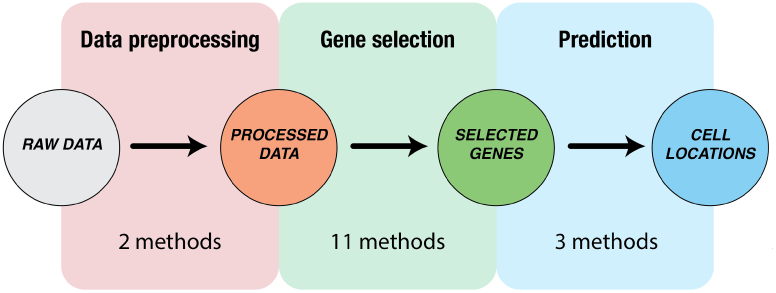
Workflow of *SCTCwhatateam*. SCTCwhatateam has three components, including data preprocessing, gene selection, and prediction. Each component has multiple methods and each pipeline is a combination of a method in each component.

### 2.2 The components of the package

In the following, we describe each of the three components of *SCTCwhatateam*. The details of the available methods for each component are described in Section 2 of the Supplementary material.

- Data pre-processing: To process data in a single cell RNA sequencing dataset, two methods, z-score Normalisation and MAGIC [8], are implemented. z-score Normalisation is used to normalise the raw data and MAGIC is used to impute missing values.
- Gene selection: To select the important genes which can be used for the prediction, we have developed 8 new methods based on the gene expression and biological information of spatial marker genes (i.e. genes in Section 1 of the Supplementary material). We also employ 3 existing unsupervised feature selection methods based on linear regression, including General Stepwise Linear Regression, Reverse Stepwise Linear Regression, and Forward Stepwise Linear Regression.
- Cell location prediction: Three methods have been implemented in the package for cell location prediction: Matthews Correlation Coefficient score (MCC) [5], Local Outlier Factor (LOF) [2], and Pearson correlation coefficient. MCC predicts cell locations based on the matching of gene expression in cells and locations. LOF is used together with MCC to eliminate outlier locations. Pearson correlation coefficient predicts cell locations by matching cells and locations on continuous sequencing data.

### 2.3 Implementation

The *R* scripts of the package are available at https://github.com/thanhbuu04/SCTCwhatateam and the vignette is illustrated in Supplement. The *Shiny* application is available at https://github.com/pvvhoang/SCTCwhatateam-ShinyApp.

## 3 Applications and conclusion

We have applied the proposed methods and workflow to predict the positions for each of the 1,297 cells in the DREAM challenge. We participated all three sub-challenges. We evaluated our methods mainly based on average distance and we used the number of bins as an additional indicator to select the methods for submissions. The three pipelines, as shown in Table 1, were submitted to the challenge and two of them were the winners of the sub-challenges [7] (see the detailed results in Supplementary material Section 3).

**Table 1.** Three pipelines submitted to the challenge. Details of the winning methods are in the Supplement.

| Pre-processing | Feature selection | #genes | Prediction | Ranking among 34 participating teams |
| --- | --- | --- | --- | --- |
| MAGIC | Seed-Based-Genes | 60 | MCC&LOF | Tied 1 |
| MAGIC | Distance-Based-Genes | 40 | MCC&LOF | 1 |
| Normalisation | Linear Regression | 20 | MCC&LOF | 3 |

To provide users the flexibility of using the *SCTCwhatateam* package, we have integrated it into a *Shiny* application. With the *Shiny* application, besides using the Drosophila dataset used by the DREAM challenge, users can upload their own dataset and then choose different methods in processing data, selecting genes, and predicting cell locations, and evaluating the predictions based on their needs.

In conclusion, we have developed an *R* package to predict cell locations and integrated it in a *Shiny* application. The effectiveness of the package is proved as we are the winners in the DREAM challenge. We hope that the tools will be useful for single cell spatial prediction research.

## Acknowledgements

Data used in this publication were generated by Prof. Dr. Nikolaus Rajewsky, Max Delbrück at the Center for Molecular Medicine, and these results were obtained as part of the DREAM Single Cell Transcriptomics Challenge project through Synapse ID (syn15665609). This work is supported by an Australian Government Research Training Program (RTP) Scholarship and the Vice Chancellor & President’s Scholarship offered by the University of South Australia.

## Funding

This work has been supported by the ARC DECRA (No: 200100200) and the Australian Research Council Discovery Grant (No: DP170101306).

## Biographical note

**Vu Viet Hoang Pham** is a PhD student at the University of South Australia (UniSA). He received his Master of Information Technology in 2017 at Deakin University, Australia. His research interests are causal inference and its applications in Bioinformatics.

**Xiaomei Li** is a PhD student at the University of South Australia (UniSA). Her research interests are causal inference and its applications in Bioinformatics.

**Buu Truong** is a visiting student at the University of South Australia (UniSA). His research interests are causal inference and its applications in Bioinformatics.

**Thin Nguyen** is a Senior Research Fellow at Applied Artificial Intelligence Institute (A2I2) at Deakin University. He received his PhD of Computer Science in 2012 at Curtin University, Australia in the area of machine learning and social media analytics. His research interests are inter-disciplinary data analytics, pattern recognition, web-scale analysis, genetics, and medicine, specifically in the areas of cancer, depression, and autism.

**Lin Liu** is an associate professor at the School of Information Technology and Mathematical Sciences, University of South Australia (UniSA). She received her bachelor and master degrees in Electronic Engineering from Xidian University, China in 1991 and 1994 respectively, and her PhD degree in computer systems engineering from UniSA in 2006. Her research interests include data mining and bioinformatics, as well as Petri nets and their applications to protocol verification and network security analysis.

**Jiuyong Li** is a professor at the School of Information Technology and Mathematical Sciences, University of South Australia. He received his PhD degree in computer science from the Griffith University, Australia (2002). His research interests are in the fields of data mining, privacy preserving and bioinformatics. His research has been supported by six prestigious Australian Research Council Discovery grants since 2005 and he has published more than 100 research papers.

**Thuc Duy Le** is a senior lecturer at the University of South Australia (UniSA). He is also an ARC DECRA fellow in Bioinformatics. He received his PhD degree in Computer Science (Bioinformatics) in 2014 at UniSA. His research interests are causal inference and its applications in bioinformatics.

## Key points

- Developing methods to predict cell locations, including methods for data preprocessing, gene slection, and location prediction
- Providing the *R* package of *SCTCwhatateam* which includes all of the methods to make the methods accessible to researchers for spatial cell reconstruction using single cell RNA-seq data
- Building a *Shiny* web-application to allow users to upload their own datasets, customise their “recipe” by selecting methods in each component of the pipeline, and download the prediction results
- Applying the proposed methods and workflow to predict the cell positions for the DREAM single cell transcriptomics challenge and being the winners in two (sub-challenges 1 and 2) out of the three sub-challenges in the DREAM challenge

## Notes

### Competing Interest Statement

The authors have declared no competing interest.

